# PARP10 promotes cellular proliferation and tumorigenesis by alleviating replication stress

**DOI:** 10.1101/337220

**Authors:** Emily M. Schleicher, Adri M. Galvan, George-Lucian Moldovan, Claudia M. Nicolae

## Abstract

During carcinogenesis, cells are exposed to increased replication stress due to replication fork arrest at sites of DNA lesions and other difficult to replicate regions. Efficient fork restart and DNA repair are important for cancer cell proliferation. We previously showed that the ADP-ribosyltransferase PARP10 interacts with the replication protein PCNA and promotes lesion bypass by recruiting specialized, non-replicative DNA polymerases. Here, we show that PARP10 is overexpressed in a large proportion of human tumors. To understand the role of PARP10 in cellular transformation, we inactivated PARP10 in HeLa cancer cells by CRISPR/Cas9-mediated gene knockout, and overexpressed it in non-transformed RPE-1 cells. We found that PARP10 promotes cellular proliferation and replication fork elongation. Mechanistically, PARP10 overexpression alleviated cellular sensitivity to replication stress by fostering the restart of stalled replication forks. Importantly, mouse xenograft studies indicated that loss of PARP10 reduces the tumorigenesis activity of HeLa cells, while its overexpression results in tumor formation by non-transformed RPE-1 cells. Our findings indicate that PARP10 promotes cellular transformation by alleviating replication stress, and suggest that targeting PARP10 may represent a novel therapeutic approach.

## INTRODUCTION

ADP-ribosylation is a post-translational modification that has recently emerged as an important regulatory factor in both DNA and cancer biology. The PARP family of ADP-ribosyltransferases encompasses 17 enzymes with a PARP (poly-ADP-ribose polymerase) catalytic domain in their C-termini (1,2). PARP1, the founding member of the family, and the closely related PARP2 and PARP3, catalyze formation of poly-ADP-ribose chains on themselves and a number of other substrates. PARP1 plays major roles in regulating DNA transcription, repair, and replication. Loss or inhibition of PARP1 results in spontaneous death of cells with Homologous Recombination (HR) DNA repair deficiency, and thus PARP1 inhibitors are used in clinical treatment of breast and ovarian tumors with BRCA1 or BRCA2 mutations (3-6)

In contrast to PARP1, which catalyzes poly-ADP-ribose chain formation, PARP10 (also known as ARTD10) and other members of the PARP family catalyze the transfer of a single ADP-ribose molecule (process known as mono-ADP-ribosylation, or MARylation) (7). In line with this, the functions of PARP10 are distinct than those of PARP1. PARP10 was originally identified as a Myc-interacting protein (8). Subsequently it has been proposed to be important for the G1/S cell cycle transition (9) as well as for caspase-dependent apoptosis (10). More recently, it was shown that PARP10 can suppress cytokine-induced activation of the NF_κ_B pathway (11), and plays roles in mitochondrial oxidation (12) and cell migration (13).

We have previously uncovered an unexpected involvement of PARP10 in DNA repair (14,15). We showed that PARP10 interacts with the replication protein PCNA, an essential polymerase co-factor (16) which recruits PARP10 to replication forks. We found that PCNA interaction is mediated by the PIP-box (PCNA-interacting peptide motif) sequence QEVVRAFY at position 834-841 in PARP10. One of the well-described roles of PCNA is promoting the stability and progress of replication machineries during stress conditions. Unrepaired DNA lesions, secondary DNA structures, repetitive elements and other non-canonical DNA structures can arrest the progression of replicative DNA polymerases (17,18). Unless efficiently restarted, stalled replication forks can disassemble, resulting in DNA strand breaks and genomic instability. One mechanism that restarts stalled replication forks is Translesion DNA Synthesis (TLS), which employs specialized polymerases able to accommodate modified DNA bases in their active sites, to bypass fork arresting structures (16,17). Upon replication fork arrest, mono-ubiquitination of PCNA at Lys164 promotes recruitment of TLS polymerases, which possess PIP and ubiquitin-interacting motifs, to re-start the stalled fork (19,20). We previously showed that PARP10 downregulation results in reduced levels of PCNA ubiquitination, impaired recruitment of the TLS polymerase Rev1 to sites of DNA damage, and sensitivity to replication arresting drugs such as hydroxyurea (HU) (14). In line with this, by employing a plasmid-based reporter of TLS activity, we showed that PARP10 is required for efficient TLS. This activity requires PCNA interaction, as TLS levels could be restored by re-expression of wildype PARP10 but not of a PARP10 variant harboring a mutation of the 8-residue PIP-box sequence (14).

During cellular transformation, increased proliferation is associated with replication stress and frequent replication fork arrest (18). Replication stress is a major barrier to oncogene-induced proliferation as it activates the DNA damage and replication stress checkpoints leading to cell cycle arrest and/or senescence (21,22). Suppression of this mechanism is an essential step in carcinogenesis. By restarting stalled replication forks, TLS suppresses DNA damage accumulation and allows completion of DNA replication, thereby enabling cellular proliferation and potentially promoting transformation (17). Because of the role of PARP10 in TLS that we previously described, we decided to investigate how PARP10 affects transformation and cancer proliferation. Here, we show that PARP10 expression promotes *in vitro* cellular proliferation and *in vivo* tumor growth by promoting replication fork stability and suppressing replication stress.

## MATERIAL AND METHODS

### Cell culture and protein techniques

Human HeLa and RPE-1 cells were grown in DMEM supplemented with 10% Fetal Calf Serum. For *PARP10* gene knockout, the commercially available PARP10 CRISPR/Cas9 KO plasmid was used (Santa Cruz Biotechnology sc-406703). Single cells were FACS-sorted into 96-well plates using a BD FACSAria II instrument. Following clonal expansion, resulting monoclonal cultures were screened by Western blot for PARP10 protein levels. For Myc-PARP10 expression, pLV-puro-TRE lentiviral constructs encoding wiltype or the ΔPARP variant spanning aminoacids 1-834 (lacking the PIP motif and PARP catalytic domain) were obtained from Cyagen. Infected cells, stably expressing the tetracycline transactivator (tTA) element, were selected by puromycin. For induction of expression, cells were grown in the presence of 2µg/ml doxycycline. Cell extracts, chromatin fractionation, and Western blot experiments were performed as previously described (14,23,24). Antibodies used for Western blot were: PARP10 (Novus NB100-2157), GAPDH (Santa Cruz Biotechnology sc-47724), Polη (Cell Signaling Technology 13848), PCNA (Cell Signaling Technology 2586), Ubiquityl-PCNA Lys164 (Cell Signaling Technology 13439).

### Functional assays

Apoptosis was quantified using the FITC Annexin V kit (Biolegend 640906). For cell cycle profiles, cells were fixed in 4% paraformaldehyde and stained with FxCycle PI reagent (Invitrogen F10797). EdU incorporation was assayed using the Click-iT Plus kit (Invitrogen C10633). For clonogenic experiments, 250 cells were seeded in 6-well plates and, two weeks later, stained with Crystal violet, or trypsinized and counted using an automated cell counter for quantification of cellular proliferation. When HU sensitivity was analyzed, cells were incubated with 0.2mM HU immediately after seeding; 72h later, media was replaced and plates were stained with Crystal violet two weeks later. For time-course proliferation experiments, 500 cells were seeded in wells of 96-well plates, and cellular viability was scored at indicated days using the CellTiterGlo reagent (Promega G7572).

### DNA Fiber Assay

Cells were incubated with 100µM CldU for 30 minutes, washed with PBS and incubated with 100µM IdU for another 45 minutes (for HeLa cells) or 90 minutes (for RPE-1 cells), either by itself, or in the presence of 0.2mM HU as indicated. Next, cells were harvested and DNA fibers were obtained using the FiberPrep kit (Genomic Vision). DNA fibers were stretched on glass slides using the FiberComb Molecular Combing instrument (Genomic Vision). Slides were incubated with primary antibodies (Abcam 6326 for detecting CIdU; BD 347580 for detecting IdU; Millipore Sigma MAB3034 for detecting DNA), washed with PBS, and incubated with Cy3, Cy5, or BV480 –coupled secondary antibodies (Abcam 6946, Abcam 6565, BD Biosciences 564879). Following mounting, slides were imaged using a Leica SP5 confocal microscope. At least 100 tracts were quantified for each sample.

### In vivo xenograft studies

Cells (either 5 million or 10 million, as indicated in the Results section) were suspended in PBS and mixed 1:1 with Matrigel Matrix (Corning 354234). Cells were then injected into both flanks of 4-6 weeks old athymic nude female mice (Charles River Laboratories stock #490). For the experiments involving cells with exogenous PARP10 expression, mice were also administered 2mg/mL doxycycline in their drinking water (supplemented with 5mg/ml sucrose) starting the day of injection.

### Statistical analyses

With the exception of the DNA fiber combing and the xenograft data, the statistical analysis performed was the TTEST (two-tailed, equal variance), using PRISM software. For the DNA fiber combing and the xenograft data, the Mann-Whitney test was performed. Statistical significance is indicated for each graph (ns = not significant, for P > 0.05; * for P < 0.05; ** for P < 0.01; *** for P < 0.001; **** for P < 0.0001).

## RESULTS

### Loss of PARP10 inhibits proliferation of HeLa cells and increases sensitivity to replication stress

To gain insights into a possible involvement of PARP10 in carcinogenesis, we mined the publicly available cBioPortal database (25) for *PARP10* gene expression and mutation in cancer samples. Strikingly, we found that, throughout a variety of cancer types and datasets, the *PARP10* gene was almost exclusively amplified, with almost no *PARP10* gene mutations or deletions being found (Supplementary Figure 1A). This pattern is very similar to that of known oncogenes (such as *MYC*), and contrasts that of known tumor suppressors, such as *BRCA2* –for which gene mutations or deletions are predominant, while only few amplifications are found (Supplementary Figure 1A). Moreover, this pattern is specific to PARP10, as the related mono-ADP-ribosyltransferase PARP14 shows a seemingly random pattern of gene amplification and deletion/mutations (Supplementary Figure 1A). Moreover, survey of cancer-specific TCGA datasets, which also include mRNA quantification, showed that up to 19% of all breast tumors and 32% of all ovarian tumors have increased PARP10 expression (gene amplification and/or mRNA upregulation) (Supplementary Figure 1B). These findings suggest that PARP10 may function as an oncogene and promote transformation.

**Figure 1.**
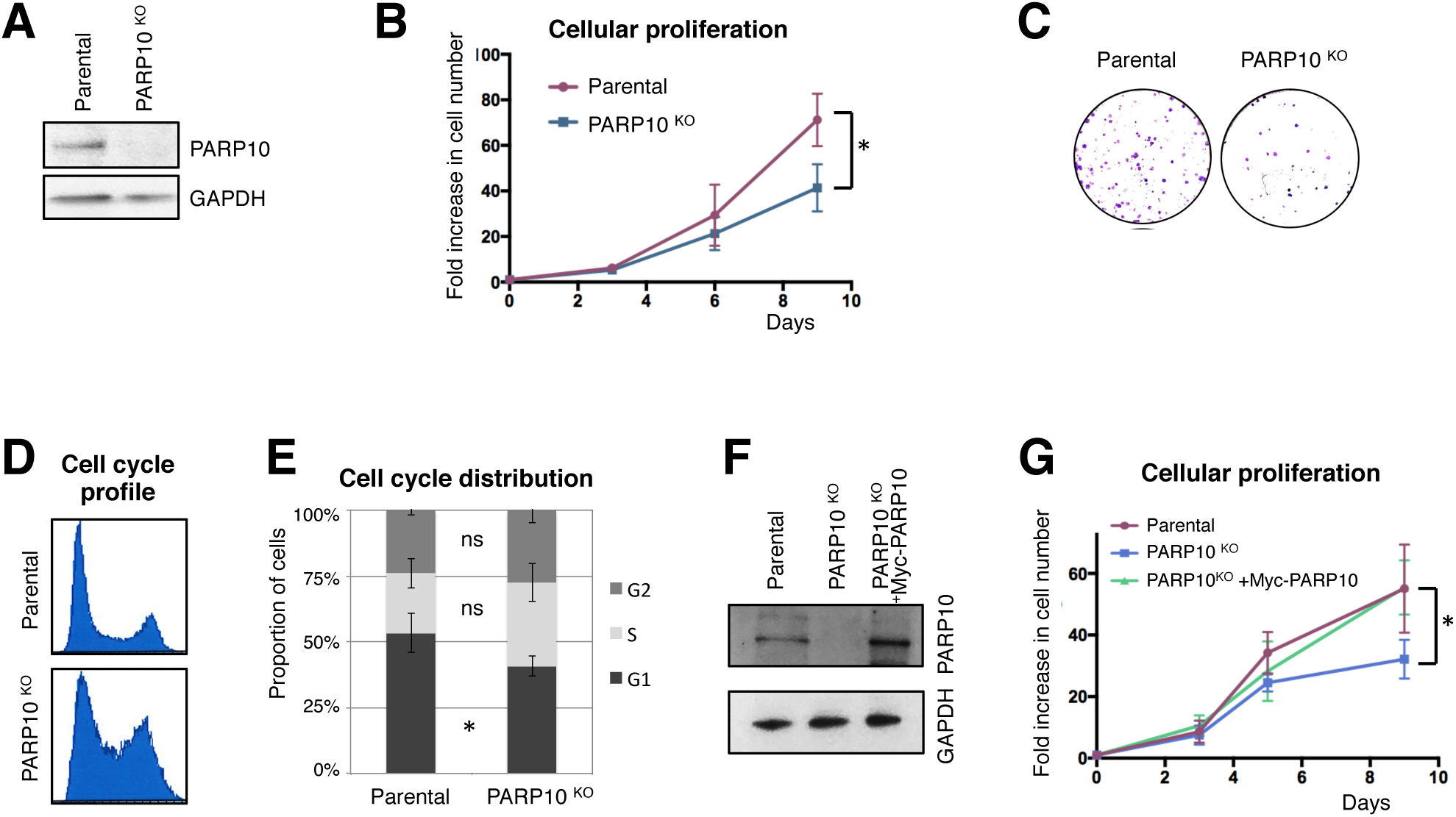
Loss of PARP10 impairs proliferation of HeLa cells. **A.** Western blot showing loss of PARP10 expression in HeLa cells with CRISPR/Cas9-mediated PARP10 knockout. **B.** PARP10-knockout HeLa cells show reduced proliferation rates. The average of 3 experiments with error bars representing standard deviations is shown. The asterisk indicates statistical significance (using the two-tailed equal variance TTEST). **C**. Representative clonogenic assay showing reduced proliferation of PARP10-knockout HeLa cells. **D**. Representative PI flow cytometry profile showing an altered cell cycle distribution in PARP10-knockout HeLa cells. **E**. Quantification of cell cycle distribution in control and PARP10-knockout HeLa cells. The average of four experiments, with error bars as standard deviations, is shown. Statistical significance was calculated using the two-tailed equal variance TTEST. **F**. Western blot showing the re-expression of PARP10, with a Myc-tag, in PARP10-knockout HeLa cells. **G**. Exogenous PARP10 expression rescues the proliferation defect of PARP10-deleted HeLa cells. The average of four experiments with error bars representing standard deviations is shown. The asterisk indicates statistical difference between the PARP10^KO^ and PARP10^KO^ + Myc-PARP10 samples.

To evaluate its impact on proliferation of cancer cells, we knocked-out PARP10 in HeLa cells using CRISPR/Cas9 technology (Figure 1A). PARP10-deleted HeLa cells showed reduced proliferation (Figure 1B, C) and an altered cell cycle profile, with increased accumulation of cells in S-phase suggestive of S-phase arrest (Figure 2D,E). Importantly, stable re-expression of wiltype Myc-tagged PARP10 from a lentiviral construct (Figure 1F) could rescue the proliferation defect of the PARP10-knockout cells (Figure 1G). These findings indicate that loss of PARP10 reduces cancer cell proliferation.

**Figure 2.**
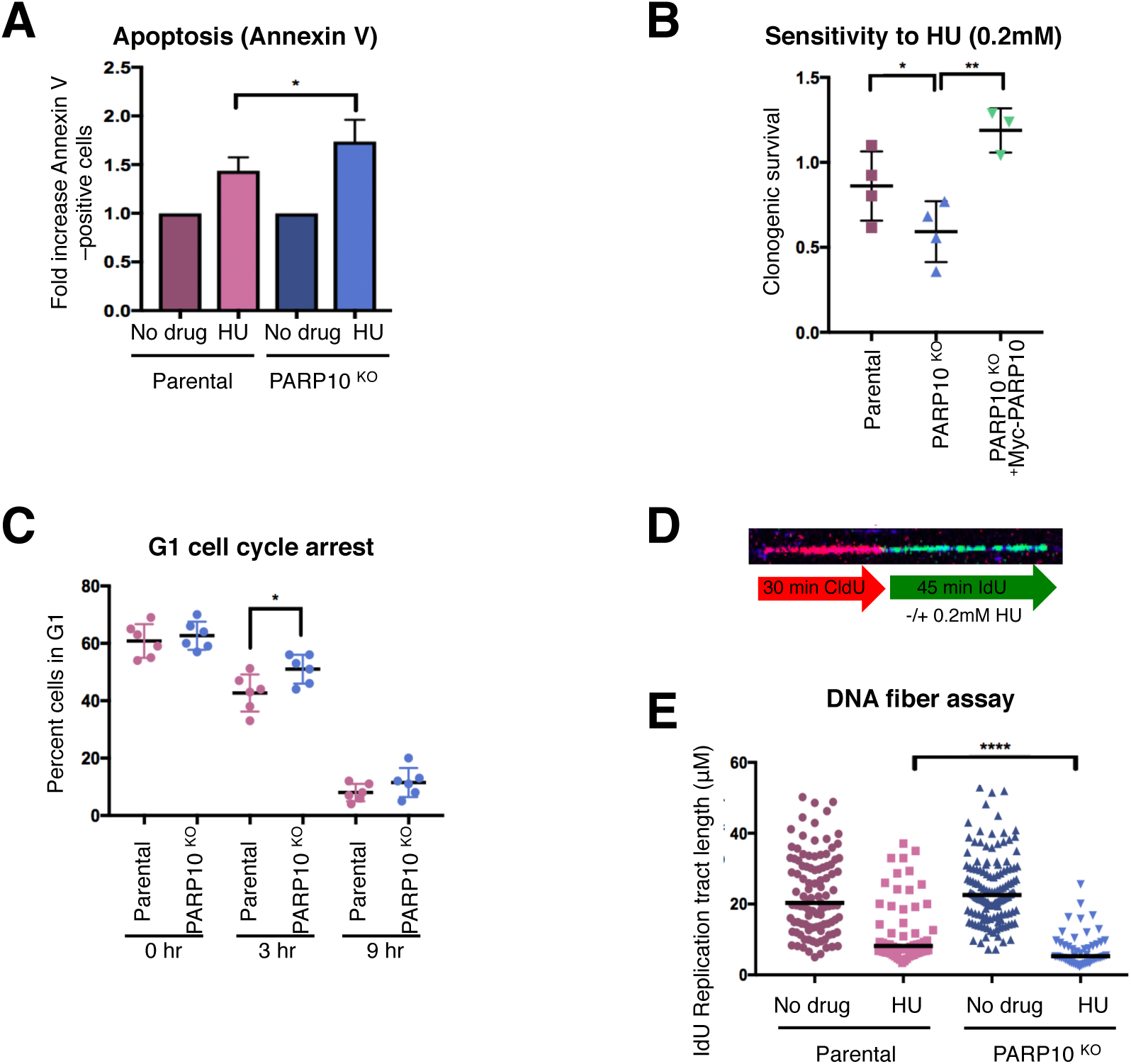
Loss of PARP10 results in sensitivity to replication stress. **A.** Annexin V apoptosis experiment showing increased apoptosis in PARP10-knockout cells following HU treatment. Cells were treated with 1mM HU for 24h. Data is shown as normalized to the control (no drug treatment) condition for each cell line. The mean of 6 experiments with error bars as standard deviations is shown. **B**. Clonogenic assay showing that PARP10-knockout cells are sensitive to HU, and re-expression of PARP10 corrects this sensitivity. Data is shown as normalized to the control (no drug treatment) condition for each cell line. The mean and standard deviation are shown. **C**. Quantification of the proportion of cells in G1 at the indicated time points after release from HU (1mM for 24h). The mean and standard deviation are shown. Representative flow cytometry histograms are shown in Supplementary Figure 2. **D**. Schematic representation of the DNA fiber combing assay condition, including a representative micrograph. **E**. DNA fiber combing assay showing reduced replication fork progression in PARP10-knockout cells upon HU exposure. Shown is the quantification of the IdU tract length, with the median values marked.

We next tested the ability of PARP10-deleted HeLa cells to handle replication stress. Exposure to hydroxyurea increased apoptosis and reduced clonogenic survival, which was restored by Myc-PARP10 re-expression (Figure 2A, B). Acute HU treatment induced G1/S cell cycle arrest to a similar extent in both control and PARP10-knockout cells. However, upon removal of drug and re-plating in fresh media, PARP10-knockout cells showed a delay in re-starting replication (Figure 2C, Supplementary Figure 2) suggesting reduced ability to recover from replication stress. Altogether, these results argue that PARP10 is important for alleviating replication stress.

Next, we investigated the progression and stability of individual replication factories at the molecular level, using the DNA fiber combing assay. Cells were grown in the presence of thymidine analogs CldU and IdU consecutively, with 0.2mM HU being added during the IdU incubation (Figure 2D). Quantification of the IdU tract length showed a significant reduction in replication tract length in PARP10-knockout cells (Figure 2E), indicating that PARP10 is required for fork elongation under replication stress conditions.

### PARP10 overexpression promotes cellular proliferation and replication fork stability

PARP10 overexpression was identified in a large proportion of tumors (Supplementary Figure 1), suggesting that PARP10 may act as an oncogene to promote transformation. To address this, we overexpressed PARP10 in the non-transformed, hTERT-immortalized human epithelial cell line RPE-1 using a stable, lentiviral-mediated, doxycycline-inducible system (Figure 3A, B). As a control, we also overexpressed a PARP10 variant lacking the PCNA-interacting PIP-box and the catalytic PARP domain (PARP10-ΔPARP, spanning residues 1-834). PARP10 overexpression resulted in increased growth rates of RPE-1 cells, while PARP10-ΔPARP overexpression did not alter proliferation (Figure 3C, D), indicating that the PCNA interaction and the catalytic activity are essential for the proliferation-promoting activity of PARP10.

**Figure 3.**
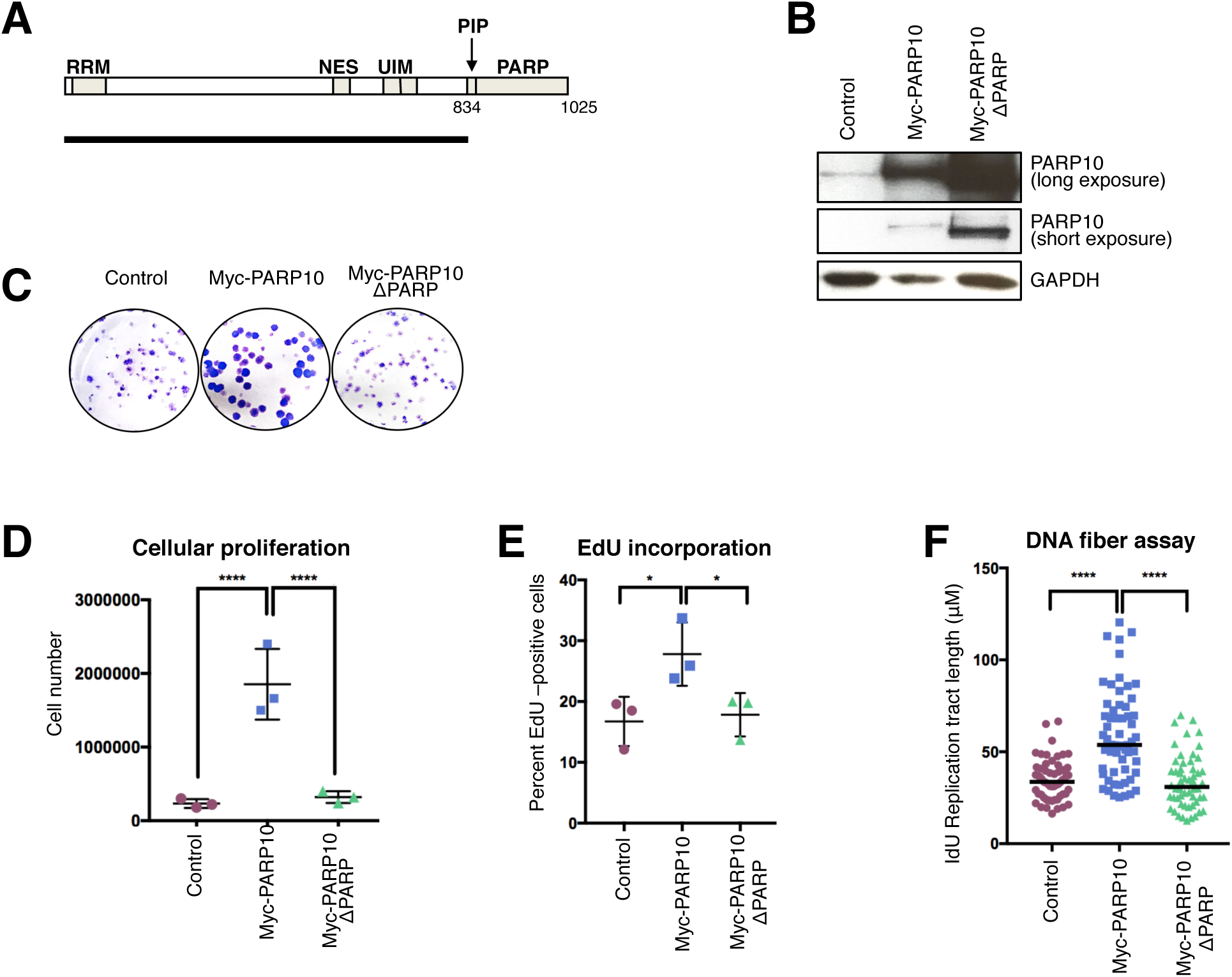
Overexpression of PARP10 promotes proliferation of non-transformed RPE-1 cells. **A.** Schematic representation of PARP10 domain organization. The black line underneath shows the length of the PARP10-ΔPARP variant (spanning residues 1-834) which lacks the PCNA-interacting PIP motif and the catalytic PARP domain. RRM: RNA recognition motif; NES: nuclear localization signal; UIM: ubiquitin interacting motifs; PIP: PCNA-interacting motif; PARP: catalytic ADP-ribosyltransferase domain. Western blot showing the overexpression of Myc-tagged PARP10 wildtype and ΔPARP in RPE-1 cells. **C**-**E**. Overexpression of wildtype, but not PARP-deleted PARP10, promotes proliferation of RPE-1 cells. Representative clonogenic assay. **D**. Quantification of cell number from clonogenic assays using CellTiterGlo reagent. The mean and standard deviation are shown. **E**. Quantification of EdU-incorporating cells. EdU was added to the media for 45 minutes prior to harvesting. The mean with standard deviation is shown. Representative flow cytometry plots are shown in Supplementary Figure 3. **F**. DNA fiber combing assay showing increased replication tracts in PARP10-overexpressing RPE-1 cells under normal (no drug treatment) conditions. Shown is the quantification of the IdU tract length, with the median values marked.

To understand the mechanistic bases for their increased proliferation, we measured DNA synthesis rates in PARP10-overexpressing RPE-1 cells. We grew cells in the presence of thymidine analog EdU for 45mins and quantified EdU incorporation using Click chemistry. PARP10-overexpressing cells showed increased EdU incorporation, while PARP10-ΔPARP overexpression did not affect EdU incorporation rates (Figure 3E, Supplementary Figure 3). Moreover, DNA fiber combing indicated that, under normal growth conditions (no drug treatment), PARP10 overexpression results in longer DNA tracts (Figure 3F), which was not the case for PARP10-ΔPARP overexpression. These findings indicate that PARP10 promotes replication fork progression in non-transformed cells.

Next, we investigated if replication fork elongation upon PARP10 overexpression is coupled with increased resistance to replication stress. Clonogenic assays indicated that PARP10 overexpressing cells, but not PARP10-ΔPARP-overexpressing cells, were resistant to HU (Figure 4A). Moreover, PARP10-overexpressing cells showed longer replication tracts in the presence of HU in the DNA fiber combing assay (Figure 4B), indicating that PARP10 overexpression promotes the ability of the replication machinery to restart stalled forks.

**Figure 4.**
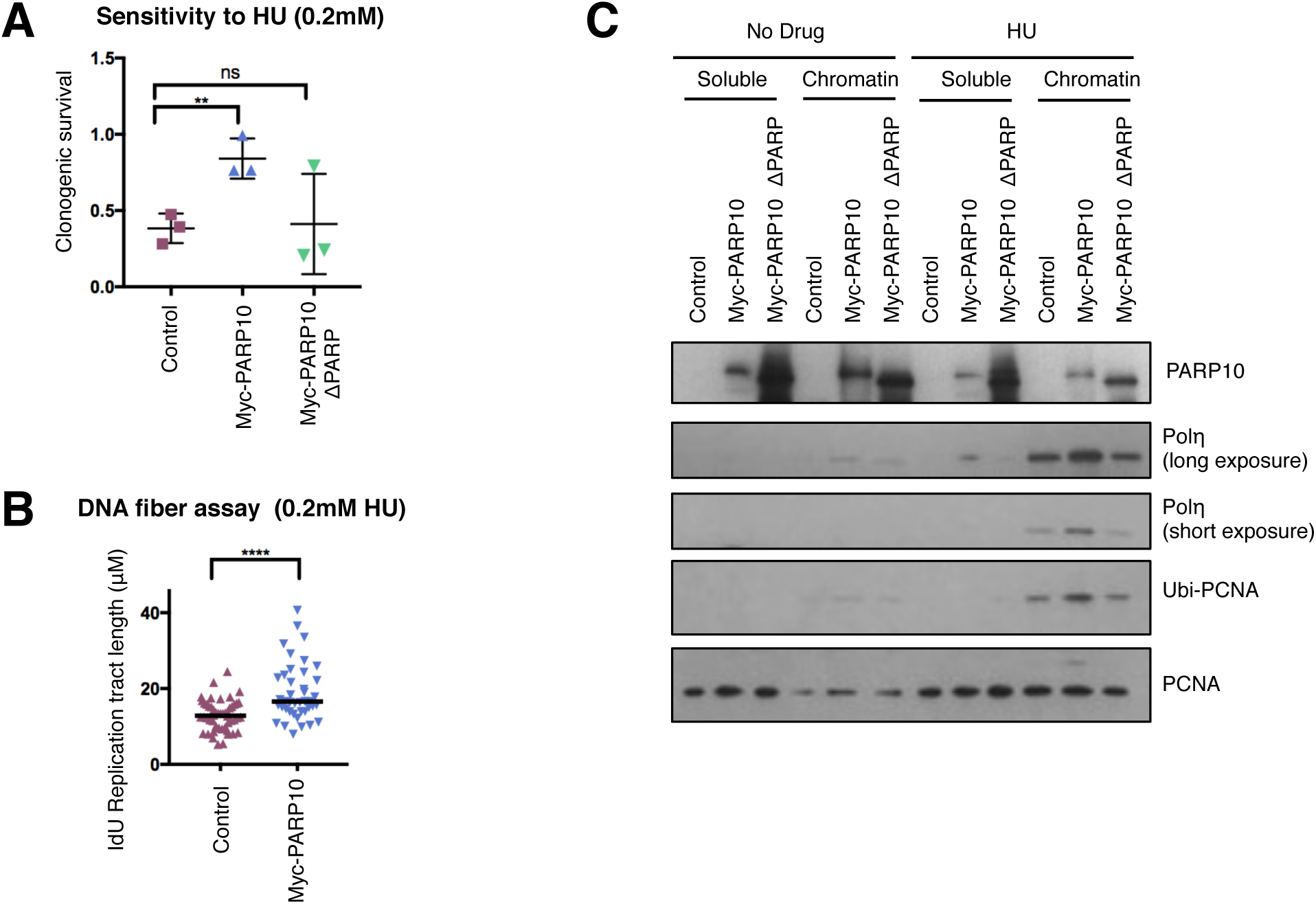
Overexpression of PARP10 in RPE-1 suppresses replication stress. **A.** Clonogenic assay showing that PARP10-overexpressing RPE-1 cells are resistant to HU. Data is shown as normalized to the control (no drug treatment) condition for each cell line. The mean and standard deviation are shown. DNA fiber combing assay showing increased replication tracts in PARP10-overexpressing RPE-1 cells under HU treatment. Shown is the quantification of the IdU tract length, with the median values marked. Chromatin fractionation experiments showing increased chromatin recruitment of TLS polymerase Polη in PARP10-overexpressing RPE-1 cells following HU treatment (2mM for 24h).

Next, we attempted to decipher how PARP10 promotes fork stability under replication stress. We previously showed that PARP10 downregulation reduces the levels of ubiquitinated PCNA (14). In line with this, overexpression of PARP10 increased PCNA ubiquitination in RPE-1 cells (Figure 4C). The TLS polymerase Polη was previously shown to be recruited to chromatin and promote DNA synthesis upon HU treatment (26). As PCNA ubiquitination targets Polη to stalled forks (20), we reasoned that PARP10 overexpression may result in increased Polη engagement to promote replication under HU conditions. Indeed, chromatin fractionation experiments showed that RPE-1 cells overexpressing wildtype, but not the ΔPARP variant, show increased chromatin loading of Polη upon HU exposure (Figure 4C). Altogether, these results indicate that PARP10 promotes PCNA ubiquitination and subsequent TLS polymerase engagement to enhance the restart of stalled replication forks and allow DNA synthesis under replication stress conditions.

### PARP10 regulates tumor growth *in vivo*

Because of the strong effect of PARP10 on *in vitro* cellular proliferation that we uncovered here, we decided to investigate the impact of PARP10 on *in vivo* tumor growth using xenograft models. First, we tested tumor formation by HeLa PARP10-knockout cells. We subcutaneously injected 5 million cells, matrigel-embedded, in each flank of athymic nude mice, and monitored tumor growth. As expected, wildtype HeLa cells generated robust tumors within four weeks from the time of injection. In contrast, PARP10-knockout cells showed severely impaired tumor formation capacity (Figure 5A, B) indicating that PARP10 is necessary for tumor growth *in vivo*. To rule out possible off-target effects generated by the CRISPR/Cas9 genome editing system in the PARP10-knockout cells, we repeated the experiment and included HeLa PARP10-knockout cells corrected by doxycycline-induced expression of Myc-tagged PARP10. For this experiment, 10 million matrigel-embedded cells were injected, and mice were administered doxycycline in their drinking water starting at the day of injection. Cells re-expressing Myc-PARP10 could form tumors similar to parental cells (Figure 5C), thus firmly establishing that PARP10 is specifically required for tumor growth *in vivo*.

**Figure 5.**
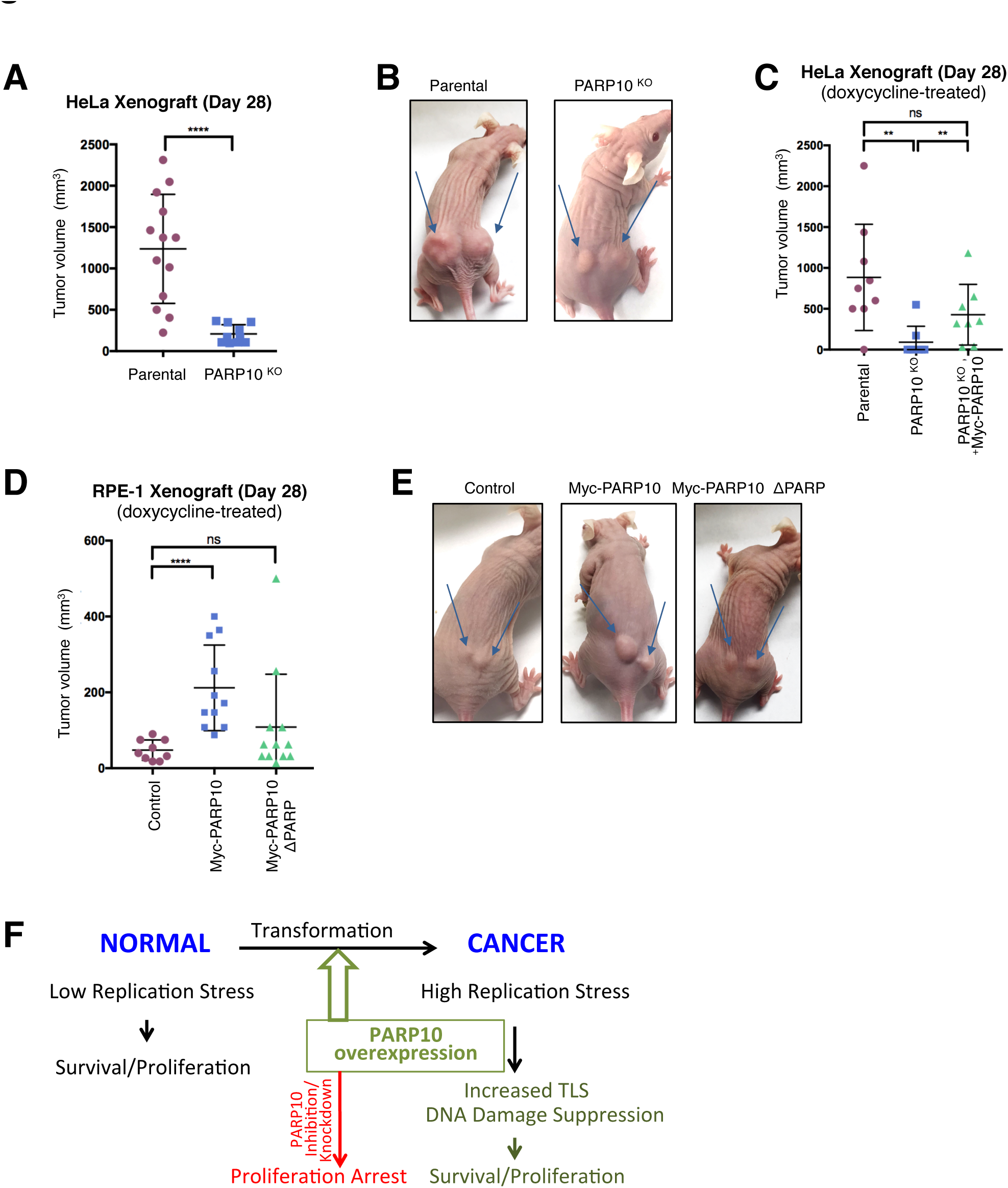
PARP10 promotes tumor growth *in vivo*. **A. B.** PARP10 deletion reduces tumor formation by HeLa cells. **A.** Quantification of tumor size, 28 days after subcutaneous injection. A total of 8 mice were used for each condition. The mean and standard deviation are shown. **B**. Representative images of tumor formation by HeLa control and PARP10-knocokout cells at day 28. **C**. Exogenous re-expression of Myc-tagged PARP10 restores tumor formation ability. Quantification of tumor size, 28 days after subcutaneous injection is shown. For each condition, 5 mice were used, which were administered doxycycline in their drinking water to induce exogenous Myc-PARP10 expression. The mean and standard deviation are shown. **D, E.** PARP10 overexpression promotes tumor formation by non-transformed RPE-1 cells. **D.** Quantification of tumor size, 28 days after subcutaneous injection. For each condition, 7 mice were used, which were administered doxycycline in their drinking water to induce exogenous Myc-PARP10 expression. The mean and standard deviation are shown. **E**. Representative images of tumor formation by RPE-1 cells at day 28. **F**. Model showing the involvement of PARP10 in carcinogenesis. PARP10 overexpression during transformation confers protection against replication stress by increasing TLS. Targeting PARP10 may reduce proliferation of cancer cells.

As PARP10 is overexpressed in a significant proportion of human cancers, suggestive of an oncogenic role (Supplementary Figure 1), we next tested tumor formation by PARP10-overexpresing non-transformed RPE-1 cells. To this end, we injected 10 million matrigel-embedded RPE-1 cells (control, PARP10-overexpressing, or PARP10-ΔPARP overexpressing) in each flank of athymic nude mice, which were also administered doxycycline. Consistent with the non-transformed status of RPE-1 cells, previous studies have shown that these cells do not induce tumor formation in immunocompromised mice (27). In line with this, in our study, control RPE-1 cells induced negligible growth at the site of injection. In contrast, PARP10-overexpressing RPE-1 cells generated noticeable tumors (Figure 5D, E), albeit much smaller than those generated by HeLa cells within the same time frame. Importantly, PARP10-ΔPARP-overexpressing RPE-1 cells lacked tumor formation ability (Figure 5D, E). Altogether, these results suggest that PARP10 has oncogene-like properties.

## DISCUSSION

We show here that PARP10 is upregulated in large number of human tumors, and its overexpression promotes cellular proliferation *in vivo* and tumor formation *in vitro* (Figures 3, 4, 5 and Supplementary Figure 1). We propose that PARP10 upregulation contributes to alleviating replication stress during cellular transformation, through increasing TLS and thus suppressing DNA damage accumulation (Figure 5F). In line with this, removal of PARP10 from cancer cells severely impairs replication stress resistance and tumor formation *in vivo* (Figures 1, 2, 5).

While significantly smaller than tumors generated by HeLa cells within the same time frame, tumor formation by PARP10-overexpressing RPE-1 cells (Figure 5) is nevertheless a significant and unexpected finding. Xenograft tumor formation by RPE-1 cells was previously established as a model for investigating and validating oncogenic mechanisms. Known oncogenes, such as Ras, have been shown to promote tumor formation in this system, which was subsequently used to confirm oncogenesis by recently discovered oncogenes such as PGBD5 (27,28). Our work thus suggests that PARP10 functions as an oncogene to promote tumor growth. In line with the concept of oncogene addiction (29), loss of PARP10 reduces proliferation of cancer cells (Figure 1). Thus, PARP10 may represent a novel target in cancer therapy.

Other roles of PARP10 have been recently described, which may mediate its impact on proliferation. One study showed that PARP10 impacts the mitochondrial oxidation process (12). While we cannot exclude an impact of this process on the tumor-promoting activity of PARP10, our data clearly show that PARP10 enhances replication fork elongation at the molecular level, thus strongly arguing that replication stress suppression through TLS engagement is a major component of the oncogenic activity of PARP10. Another recent paper described a role for PARP10 in cell motility through regulating Aurora A activity (13). Interestingly, the authors also created PARP10-knockout HeLa cells and found increased cell motility *in vitro,* and increased metastasis *in vivo* (using an experimental approach in which they injected the HeLa cells in the tail vein of Balb/C mice and measured lung metastases). As the experimental setup is different from our study, we believe that these are likely to be two separate functions of PARP10. Our data clearly show that PARP10 knockout results in reduced proliferation *in vivo* and tumor formation *in vitro*, and both phenotypes can be rescued by exogenous re-expression of PARP10. Moreover, we show that this correlates with replication fork stability defects and replication stress hypersensitivity. Thus, regardless of any effects of PARP10 on cell motility, our data strongly support a role for PARP10 and its interaction with PCNA in alleviating replication stress and promoting proliferation.

Our work shows that PARP10 overexpression enhances engagement of TLS polymerases to promote DNA synthesis under endogenous and exogenous replication stress. Indeed, PARP10 overexpressing cells show longer replication tracts under both control (no drug) and HU treatment (Figures 3, 4). While our DNA fiber combing assays cannot differentiate between increased fork speed versus more efficient restart of stalled forks, our results showing engagement of Polη suggest that PARP10 overexpression acts through fork restart. Thus, our studies provide additional support for the model that, by alleviating replication stress, TLS promotes cellular transformation (17). Coupled with our previous study showing that PARP10 downregulation reduces mutation rates (14), these findings also suggest that overexpression of PARP10 is potentially associated with increased mutagenesis. This induction of genomic instability is likely to contribute to the oncogenic activity of PARP10.

At this time, it is still unclear exactly how PARP10 promotes TLS. Its effect of promoting PCNA ubiquitination (see our previous work (14) and Figure 4), suggests that PARP10 activity may directly increase the enzymatic process of PCNA ubiquitination, either by making PCNA a better substrate or by enhancing the activity of ubiquitin ligases such as RAD18 towards PCNA (16). Alternatively, PARP10 may act downstream of this modification by stabilizing it against de-ubiquitination by USP1. Regardless of the exact mechanism, our results using the PARP10-ΔPARP mutant indicate that PCNA binding and/or the catalytic activity of PARP10 are required for this activity. In line with this, in preliminary studies (not shown) we observed that PARP10 is able to ADP-ribosylate PCNA *in vitro*. We speculate that this modification is in turn allowing PCNA ubiquitination levels to build up, thus increasing TLS polymerase recruitment to promote DNA synthesis under replication stress conditions.

## ACKNOWLEDGEMENTS

We would like to thank Dipanjan Chowdhury and Katherine Choe for the RPE-1 cells, and the Penn State College of Medicine Flow Cytometry and Imaging cores. This project is funded, in part, under a grant with the Pennsylvania Department of Health using Tobacco CURE Funds. The Department specifically disclaims responsibility for any analyses, interpretations or conclusions.

## Legends to Supplementary Figures

**Supplementary Figure 1.**
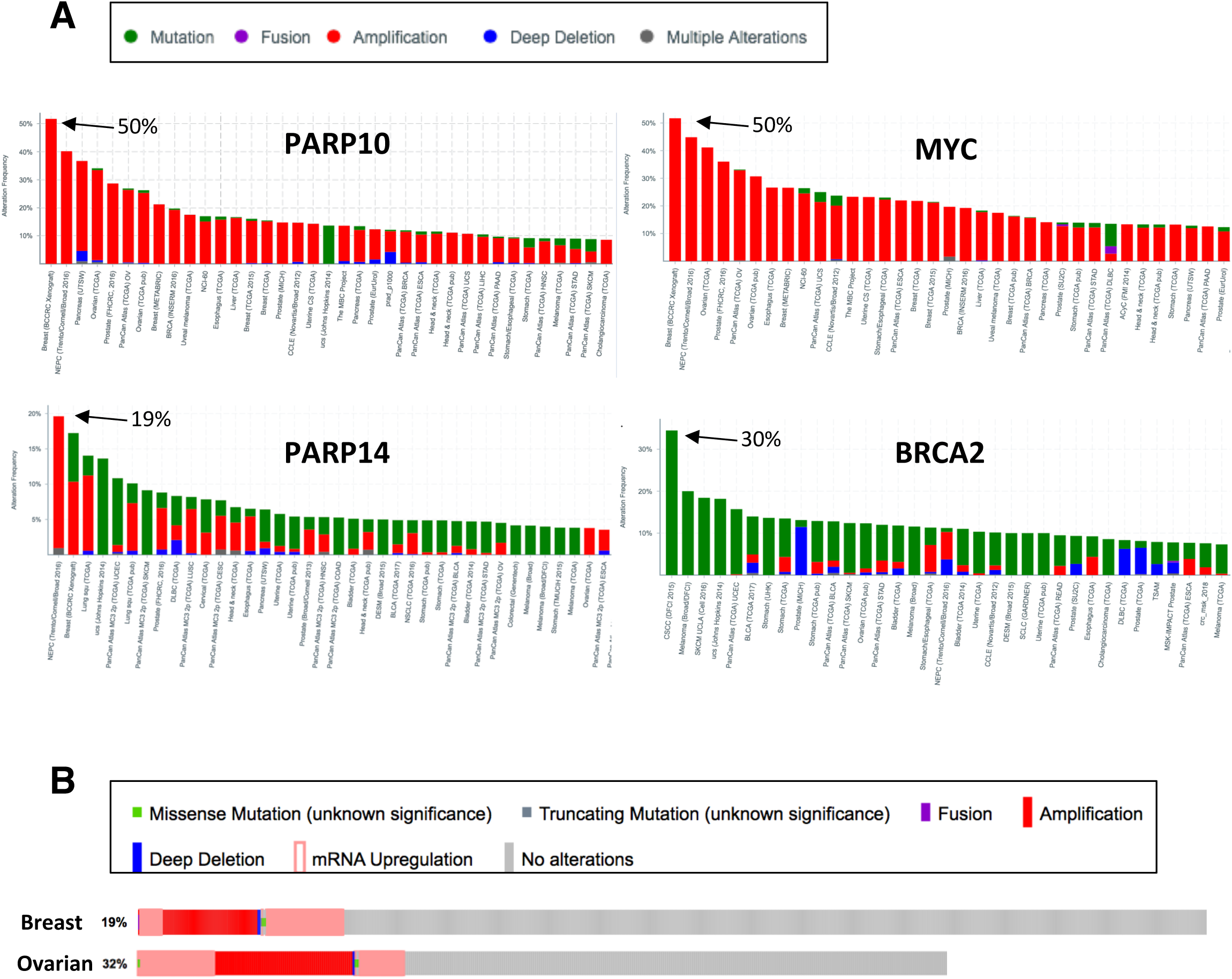
cBioPortal database mining showing PARP10 overexpression/amplification in cancer. **A**. Representation of gene mutations (green) or amplification (red) in various cancers. PARP10 shows amplifications exclusively, similar to known oncogene Myc. In contrast, known tumor suppressor BRCA2 shows mutations exclusively, while a related PARP, namely PARP14, shows a mixture of amplification and mutations –expected for genes not involved in transformation. **B**. Analyses of TCGA PanCancer samples shows that 19% of breast cancer and 32% of ovarian cancer samples investigated show PARP10 gene amplification or mRNA overexpression, and none show deletion or downregulation.

**Supplementary Figure 2.**
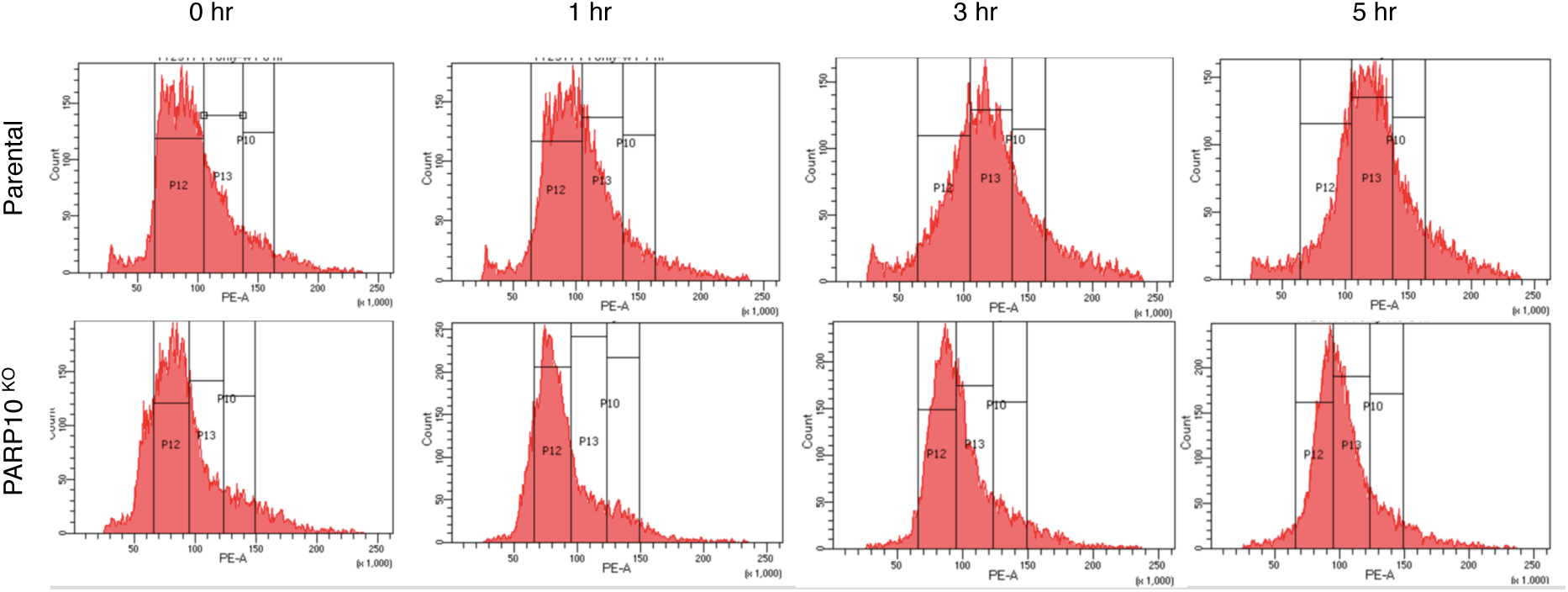
Representative flow cytometry PI histogram showing the cell cycle profile of parental and PARP10-knockout HeLa cells upon release from HU arrest.

**Supplementary Figure 3.**
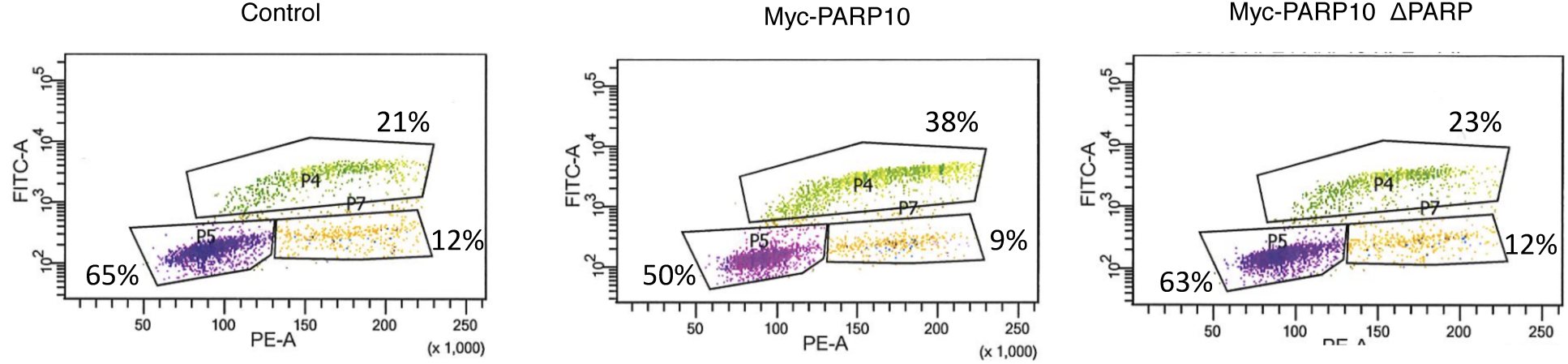
Representative flow cytometry plot showing increased EdU incorporation by PARP10-overexpressing RPE-1 cells.

## REFERENCES

1. Gibson, B.A. and Kraus, W.L. (2012) New insights into the molecular and cellular functions of poly(ADP-ribose) and PARPs. Nat Rev Mol Cell Biol, 13, 411–424.

2. Kalisch, T., Ame, J.C., Dantzer, F. and Schreiber, V. (2012) New readers and interpretations of poly(ADP-ribosyl)ation. Trends Biochem Sci, 37, 381–390.

3. Bryant, H.E., Schultz, N., Thomas, H.D., Parker, K.M., Flower, D., Lopez, E., Kyle, S., Meuth, M., Curtin, N.J. and Helleday, T. (2005) Specific killing of BRCA2-deficient tumours with inhibitors of poly(ADP-ribose) polymerase. Nature, 434, 913–917.

4. Farmer, H., McCabe, N., Lord, C.J., Tutt, A.N., Johnson, D.A., Richardson, T.B., Santarosa, M., Dillon, K.J., Hickson, I., Knights, C. et al. (2005) Targeting the DNA repair defect in BRCA mutant cells as a therapeutic strategy. Nature, 434, 917–921.

5. Fong, P.C., Boss, D.S., Yap, T.A., Tutt, A., Wu, P., Mergui-Roelvink, M., Mortimer, P., Swaisland, H., Lau, A., O’Connor, M.J. et al. (2009) Inhibition of poly(ADP-ribose) polymerase in tumors from BRCA mutation carriers. N Engl J Med, 361, 123–134.

6. Murai, J. (2017) Targeting DNA repair and replication stress in the treatment of ovarian cancer. Int J Clin Oncol, 22, 619–628.

7. Kleine, H., Poreba, E., Lesniewicz, K., Hassa, P.O., Hottiger, M.O., Litchfield, D.W., Shilton, B.H. and Luscher, B. (2008) Substrate-assisted catalysis by PARP10 limits its activity to mono-ADP-ribosylation. Mol Cell, 32, 57–69.

8. Yu, M., Schreek, S., Cerni, C., Schamberger, C., Lesniewicz, K., Poreba, E., Vervoorts, J., Walsemann, G., Grotzinger, J., Kremmer, E. et al. (2005) PARP-10, a novel Myc-interacting protein with poly(ADP-ribose) polymerase activity, inhibits transformation. Oncogene, 24, 1982– 1993.

9. Chou, H.Y., Chou, H.T. and Lee, S.C. (2006) CDK-dependent activation of poly(ADP-ribose) polymerase member 10 (PARP10). J Biol Chem, 281, 15201–15207.

10. Herzog, N., Hartkamp, J.D., Verheugd, P., Treude, F., Forst, A.H., Feijs, K.L., Lippok, B.E., Kremmer, E., Kleine, H. and Luscher, B. (2013) Caspase-dependent cleavage of the mono-ADP-ribosyltransferase ARTD10 interferes with its pro-apoptotic function. FEBS J, 280, 1330–1343.

11. Verheugd, P., Forst, A.H., Milke, L., Herzog, N., Feijs, K.L., Kremmer, E., Kleine, H. and Luscher, B. (2013) Regulation of NF-kappaB signalling by the mono-ADP-ribosyltransferase ARTD10. Nat Commun, 4, 1683.

12. Marton, J., Fodor, T., Nagy, L., Vida, A., Kis, G., Brunyanszki, A., Antal, M., Luscher, B. and Bai, P. (2018) PARP10 (ARTD10) modulates mitochondrial function. PLoS One, 13, e0187789.

13. Zhao, Y., Hu, X., Wei, L., Song, D., Wang, J., You, L., Saiyin, H., Li, Z., Yu, W., Yu, L. et al. (2018) PARP10 suppresses tumor metastasis through regulation of Aurora A activity. Oncogene.

14. Nicolae, C.M., Aho, E.R., Vlahos, A.H., Choe, K.N., De, S., Karras, G.I. and Moldovan, G.L. (2014) The ADP-ribosyltransferase PARP10/ARTD10 interacts with proliferating cell nuclear antigen (PCNA) and is required for DNA damage tolerance. J Biol Chem, 289, 13627–13637.

15. Shahrour, M.A., Nicolae, C.M., Edvardson, S., Ashhab, M., Galvan, A.M., Constantin, D., Abu-Libdeh, B., Moldovan, G.L. and Elpeleg, O. (2016) PARP10 deficiency manifests by severe developmental delay and DNA repair defect. Neurogenetics, 17, 227–232.

16. Choe, K.N. and Moldovan, G.L. (2017) Forging Ahead through Darkness: PCNA, Still the Principal Conductor at the Replication Fork. Mol Cell, 65, 380–392.

17. Zafar, M.K. and Eoff, R.L. (2017) Translesion DNA Synthesis in Cancer: Molecular Mechanisms and Therapeutic Opportunities. Chem Res Toxicol, 30, 1942–1955.

18. Zeman, M.K. and Cimprich, K.A. (2014) Causes and consequences of replication stress. Nat Cell Biol, 16, 2–9.

19. Hoege, C., Pfander, B., Moldovan, G.L., Pyrowolakis, G. and Jentsch, S. (2002) RAD6-dependent DNA repair is linked to modification of PCNA by ubiquitin and SUMO. Nature, 419, 135–141.

20. Kannouche, P.L., Wing, J. and Lehmann, A.R. (2004) Interaction of human DNA polymerase eta with monoubiquitinated PCNA: a possible mechanism for the polymerase switch in response to DNA damage. Mol Cell, 14, 491–500.

21. Bartkova, J., Rezaei, N., Liontos, M., Karakaidos, P., Kletsas, D., Issaeva, N., Vassiliou, L.V., Kolettas, E., Niforou, K., Zoumpourlis, V.C. et al. (2006) Oncogene-induced senescence is part of the tumorigenesis barrier imposed by DNA damage checkpoints. Nature, 444, 633–637.

22. Gorgoulis, V.G., Vassiliou, L.V., Karakaidos, P., Zacharatos, P., Kotsinas, A., Liloglou, T., Venere, M., Ditullio, R.A. Jr., Kastrinakis, N.G., Levy, B. et al. (2005) Activation of the DNA damage checkpoint and genomic instability in human precancerous lesions. Nature, 434, 907– 913.

23. Choe, K.N., Nicolae, C.M., Constantin, D., Imamura Kawasawa, Y., Delgado-Diaz, M.R., De, S., Freire, R., Smits, V.A. and Moldovan, G.L. (2016) HUWE1 interacts with PCNA to alleviate replication stress. EMBO Rep, 17, 874–886.

24. Nicolae, C.M., Aho, E.R., Choe, K.N., Constantin, D., Hu, H.J., Lee, D., Myung, K. and Moldovan, G.L. (2015) A novel role for the mono-ADP-ribosyltransferase PARP14/ARTD8 in promoting homologous recombination and protecting against replication stress. Nucleic Acids Res, 43, 3143–3153.

25. Gao, J., Aksoy, B.A., Dogrusoz, U., Dresdner, G., Gross, B., Sumer, S.O., Sun, Y., Jacobsen, A., Sinha, R., Larsson, E. et al. (2013) Integrative analysis of complex cancer genomics and clinical profiles using the cBioPortal. Sci Signal, 6, pl1.

26. de Feraudy, S., Limoli, C.L., Giedzinski, E., Karentz, D., Marti, T.M., Feeney, L. and Cleaver, J.E. (2007) Pol eta is required for DNA replication during nucleotide deprivation by hydroxyurea. Oncogene, 26, 5713–5721.

27. Henssen, A.G., Koche, R., Zhuang, J., Jiang, E., Reed, C., Eisenberg, A., Still, E., MacArthur, I.C., Rodriguez-Fos, E., Gonzalez, S. et al. (2017) PGBD5 promotes site-specific oncogenic mutations in human tumors. Nat Genet, 49, 1005–1014.

28. Hahn, W.C., Counter, C.M., Lundberg, A.S., Beijersbergen, R.L., Brooks, M.W. and Weinberg, R.A. (1999) Creation of human tumour cells with defined genetic elements. Nature, 400, 464–468.

29. Weinstein, I.B. and Joe, A.K. (2006) Mechanisms of disease: Oncogene addiction-a rationale for molecular targeting in cancer therapy. Nat Clin Pract Oncol, 3, 448–457.

